# Controlling spatio-temporal sequences of neural activity by local synaptic changes

**DOI:** 10.1101/2025.07.24.666534

**Authors:** Hauke O. Wernecke, Andrew B. Lehr, Arvind Kumar

## Abstract

The neural basis of behavior is believed to consist of sequential patterns of neural activity in the relevant brain regions. Behavioral flexibility also requires neural circuit mechanisms that support dynamic control of sequential activity. However, mechanisms to control and reconfigure sequential activity have received little attention.

Here, we show that recurrently connected networks with heterogeneous connectivity and a smooth spatial in-degree landscape (which may arise due to asymmetric neuron morphologies) provide a robust mechanism to evoke and control sequential activity. By modulating the synaptic strength of only a few neurons in local neighborhoods, we uncovered high-impact locations which can start, stop, extend, gate, and redirect sequences. Interestingly, high-impact locations coincide with mid in-degree regions. We demonstrate that these motifs can flexibly reconfigure sequential activity, and hence, provide a framework for fast and flexible computations on behavioral time scales, while the individual parts of the pathways remain rigid and reliable.

## Introduction

Meaningful behavior relies on ordered sequences of thought and action (Lashley 1951), which can be reconfigured in a task- or context-dependent manner. In the brain, this requires that neural circuits can not only generate neural activity sequences, but also that sequences can be flexibly modulated, gated, merged, and split. Consistent with this idea, sequential activity has been observed in a variety of tasks such as motor control (Lindén et al. 2022; Gallego et al. 2018; Eichenlaub et al. 2020), decision-making (Harvey, Coen, and Tank 2012), episodic memory encoding in the hippocampus (O’Keefe 1976; Dragoi and Buzsaki 2006; Foster and Wilson 2006; Diba and Buzsáki 2007; Pastalkova et al. 2008; Buzsáki and Tingley 2018), olfactory processing (Friedrich and Laurent 2001), and birdsong generation (Hahnloser, Kozhevnikov, and Fee 2002). Crucially, networks in the brain are able to generate neural activity sequences without sequential input, and the emergence of a specific neural activity sequence depends on the context or task (Hahnloser, Kozhevnikov, and Fee 2002; Pastalkova et al. 2008; Jin, Fujii, and Graybiel 2009; Harvey, Coen, and Tank 2012; Modi, Dhawale, and Bhalla 2014).

The emergence of sequences requires the formation of feedforward pathways (functional or anatomical) in a recurrent network. In some cases, given the finite size of networks, even random connectivity can give rise to feedforward pathways in an otherwise random network (Vogels and Larry F Abbott 2005; Lindén et al. 2022). Attractor dynamics combined with spike frequency adaptation or synaptic depression, supervised learning (i.e., echo-state networks), and unsupervised learning can all facilitate the formation of feedforward networks (Rabinovich et al. 2006; Rajan, Harvey, and Tank 2016; Rokni and Sompolinsky 2012; Zhang 1996). These mechanisms assume that the (initial) network connectivity is homogeneous and ‘symmetry of connectivity’ has to be broken by dynamics, neuron properties, or learning. Neuron morphology can also form the basis for breaking the symmetry of connectivity and give rise to the emergence of sequential activity. Axonal projections and dendritic branches extend out asymmetrically in a preferred direction (Jiang et al. 2015; Mohan et al. 2015; Xiangmin Xu et al. 2016; Peng et al. 2021; Weiler et al. 2022; Narayanan et al. 2015). Furthermore, recent data suggest that in mouse sensory cortex, nearby neurons project preferentially in similar directions, aligned with the propagation of traveling waves of activity (Ye et al. 2023). (Spreizer, Aertsen, and Kumar 2019) showed that if a distance-dependent connectivity rule is modified to account for neurons’ asymmetric shapes, feedforward pathways emerge naturally and provide the anatomical substrate for neuronal activity sequences.

In contrast to the mechanisms underlying the emergence of sequences, mechanisms to control and reconfigure sequential activity have received little attention. Synaptic plasticity would be too slow to reconfigure interactions among neural sequences on behavioral time scales. Mechanisms based on attractors with adaptation (in synapses or neurons) would rely on continuous external steering. Thus, the problem of flexible control of neural activity sequences on behavioral time scales remains poorly understood.

The model proposed by Spreizer, Aertsen, and Kumar 2019 appears at first glance too hardwired to suggest any mechanism for controlling neural activity sequences. However, here we show that spatially anisotropic connectivity also gives rise to certain circuit motifs that can be used to control sequential dynamics. In particular, in networks with spatially correlated anisotropy, neurons’ in-degree becomes heterogeneous and local regions of low, medium, or high in-degree emerge (Fig. 1). We show that in such networks, even small changes in the local connectivity can introduce large changes in the dynamics and interaction among sequences. Somewhat counterintuitively, a subset of locations with mid-range in-degree have the biggest impact on the sequence dynamics.

**Figure 1.**
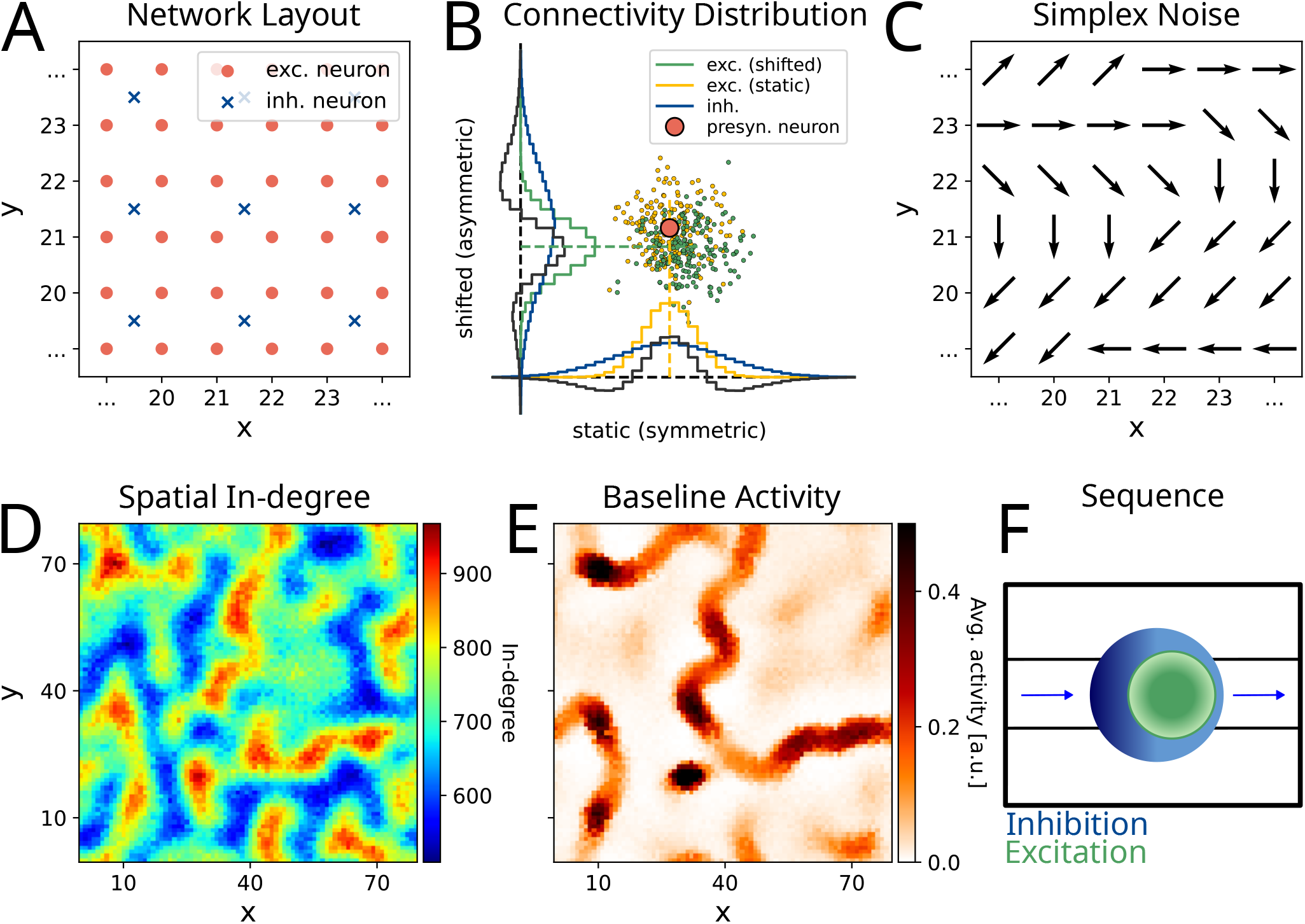
Network architecture and dynamics **(A)** Network layout: red dots represent the location of excitatory neurons, and blue crosses those of inhibitory ones. The network has a toroidal structure to avoid boundary effects. **(B)** Connectivity k ernel: The pre-synaptic neuron (red dot) shifts its targets (yellow) towards a preferred direction *ϕ* (here: bottom right), resulting in new post-synaptic targets (green). A histogram of the targets (x-, and y-axis) resembles then an asymmetric Mexican hat distribution (blue represents the inhibitory connectivity kernel, gray the combination of excitatory and inhibitory kernels). **(C)** Schematics of preferred directions *ϕ*’s which were chosen using a gradient noise algorithm (Simplex noise (Perlin 2001)). **(D)** The in-degree of the excitatory neurons reveals high- and low in-degree regions. **(E)** Average activation of the excitatory neurons. As expected higher activation occurs on regions with high in-degree. **(F)** Schematic of how activation of a small group of neurons created an excitatory center (green) and inhibitory surround (blue). Notice the asymmetry of the two regions which forms the basis of moving sequential activity. Darker colors represent stronger excitation and inhibition, respectively.

We systematically uncover motifs where small modulation of local connectivity can modulate the neural activity, i.e., start, stop, extend, gate, and select sequences. In our model, motifs are found in medium in-degree regions that act as ‘switches’ of different kinds to form the basis of rapid reconfiguration of sequential activity in the network. The existence of these motifs is dependent on the geometry of the spatially correlated anisotropy of connectivity. Thus, our work provides insights into how the spatial structure of network connectivity could be exploited for computations.

Neuromodulators are released in a patchy manner (Patriarchi et al. 2018; Sun et al. 2018; Glaeser-Khan et al. 2024), and can modulate neural excitability and synaptic strength on a behavioral time scale (Nadim and Bucher 2014; Colangelo et al. 2019). Our results provide a putative mechanism by which neuromodulation can introduce qualitative changes in network dynamics and a repertoire of intrinsic sequences, thereby contributing to behavioral flexibility and learning new skills.

## Results

We are interested in understanding how spatio-temporal activity sequences (STAS) in neuronal networks can be modified or controlled. To investigate this, we used the model proposed by Spreizer, Aertsen, and Kumar (2019) to generate STAS. For the control of STAS dynamics, we studied the effect of changing a small fraction of synapses in a small neighborhood mimicking local release of a neuromodulator (NM), which could enhance or reduce synaptic strength, and characterized the lifetime and number of unique STAS.

### Model of spatio-temporal sequences

Typically, in networks with distance-dependent connectivity, neurons connect in all directions with equal probability. When the spatial extent of inhibitory neurons exceeds that of excitatory neurons, the network dynamics exhibits a multiple bump solution (Amari 1977; Gozel and Doiron 2024). Spreizer, Aertsen, and Kumar (2019) showed that when either excitatory or inhibitory neurons project in a spatially asymmetric manner, such that 2-5% of their targets lie in a preferential direction *ϕ*, and *ϕ*’s of neighboring neurons are similar (Fig. 1A), the multiple bump solution becomes unstable and STAS emerge. This modification in the connectivity rule results in an asymmetric Mexican hat-type connectivity kernel (Fig. 1B).

Here we implemented the connectivity rule proposed by Spreizer, Aertsen, and Kumar (2019) in a network where each neuron was modeled as a firing rate-based model (see Methods, Fig. 1A-C). Spreizer, Aertsen, and Kumar (2019) argued that when neurons had a preferred direction for their projections, they formed anatomical chains that composed the basis of temporal sequences in the network. Moreover, the spatial correlation among preferred directions imposed constraints on the number and length of sequences. Besides this, we also found that even though all neurons had the same out-degree, the in-degree of neurons showed a wide distribution and a spatial structure (Fig. 1D). STAS dynamics unfolded in regions of high and mid in-degree (red and green regions), which we refer to as ‘pathways’. Because in our model inhibitory neurons projected in an isotropic manner, this spatial landscape of excitatory in-degree directly implied a landscape of high and low excitation-inhibition (EI) ratios.

It has already been described that structural variability in the connectivity can lead to an inhomogeneous distribution of EI ratios (Landau et al. 2016). Here we show that the variable EI ratio may also have a spatial structure. In our network, EI ratio variability did not lead to network-level instabilities; therefore, we did not seek to balance the individual neurons. Instead, we explore how the spatial distribution of EI ratios can be leveraged to control the STAS dynamics.

We injected uncorrelated Gaussian white noise to all the neurons to initiate the dynamics. STAS then formed in high in-degree regions and traversed along neurons with aligned preferred directions (Fig. 1E). Given the asymmetric connectivity of excitatory neurons, traveling excitatory activity was accompanied by an inhibitory ‘shadow’ (Fig. 1F). This inhibitory shadow formed the basis of interaction among STAS. Due to the long-range inhibition, multiple STAS might compete along the same pathway, or even across neighboring pathways. Overall, multiple factors like the input drive, the connectivity, as well as the spatial structure of the preferred projection directions of the neurons (*ϕ*) could affect the network dynamics.

### Effect of local change in synaptic connectivity on sequence dynamics

As the activity unfolded along high in-degree regions, the spatial structure of the EI ratios suggested that network response to a spatial external stimulus or transient perturbation should depend on its location (Fig. 1D, E). Perhaps more interestingly, the spatial inhomogeneities in the EI balance also suggested that the effect of changing synaptic weights in local patches of the network would have a bigger effect on the network activity than non-local changes. That is, changes in synaptic weights within a local neighborhood (as might happen during local release of neuromodulators) could be used as a way to control or shape the dynamics of STAS in a task- or context-dependent manner.

To illustrate and characterize this effect, we randomly selected 50 (out of 113) excitatory neurons from a circular region with a radius of 6 grid points (or neurons). For these neurons, the strengths of incoming synapses were then changed by ±20%. For the quantification of the effect of such a perturbation in the network, we measured the number of STAS and their duration in the network (see Methods). We observed that changing incoming excitatory-to-excitatory synapses of as few as 50 neurons (only 0.78% of the entire population) increased the number and decreased the duration of STAS in some cases, while in other cases, the opposite was observed (Fig. 2A, left). A similar effect was observed when we decreased the synaptic strength of incoming synapses of a local group of neurons (Fig. 2A, right). It is worth mentioning that some landscapes evoked a local static activation bump besides sequential activity in the remaining network. Similarly, a (positive) modulatory patch rarely lead to the formation of such static activity bump. We excluded those networks and patches from our analysis.

**Figure 2.**
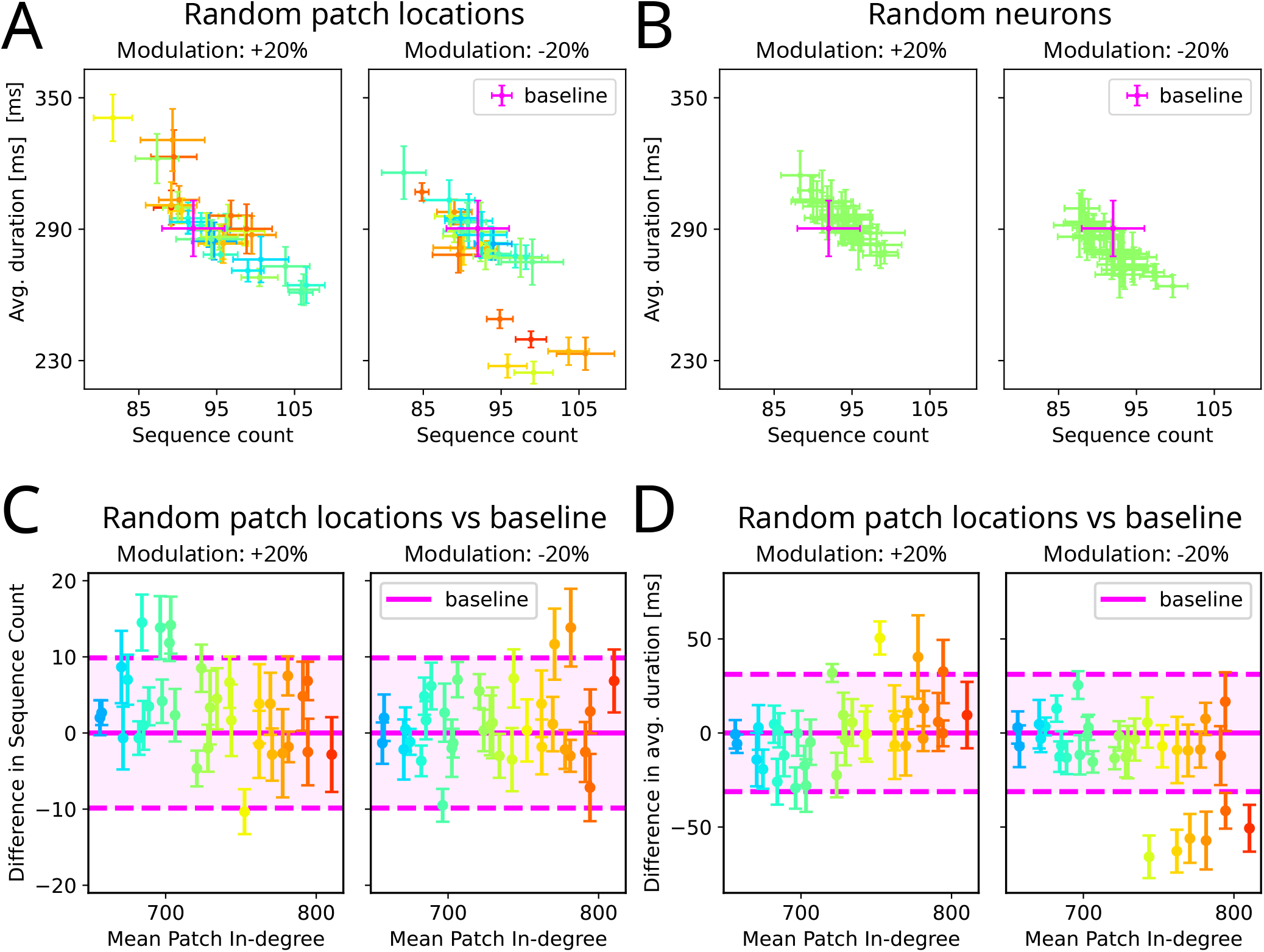
Effect of local and non-local changes in synaptic connectivity on the number of sequences and their duration. **(A)** Random spatial patch locations: In each location neurons from a small neighborhood were chosen and their connectivity was either increased or decreased by 20%. Such a modulation of connectivity resulted in large changes in the number and/or duration of STAS (error bars are the sample standard error of the mean (SEM) across random seeds of input noise). Here, results from 33 randomly chosen locations are shown. **(B)** Non-spatial NM release: In each trial 50 neurons were randomly chosen from the whole network and their connectivity was either increased or decreased by 20%. Such distributed stimulation only slightly affected the STAS. **(C)** Difference in sequence count of STAS compared to baseline over the mean in-degree of the patch location. The shaded area indicate the SEM across the baseline simulations. Locations as in A. **(D)** Same as in the panel **C**, but the difference in average duration. Locations as in A.

As a control, we also tested the effect of increasing or decreasing the strength of incoming synapses of 50 randomly chosen neurons from throughout the network. This synaptic connectivity change did not strongly alter the STAS count and duration (Fig. 2B).

Spatially localized changes in synaptic strengths resulted in a wide range of responses. Part of this variance can be explained by the modulation sign and by the location or the in-degree of the stimulated neurons (Fig. 2C, D). The strongest effects were observed when the synapses were altered in mid in-degree regions. Positive modulation seemed to be potent in increasing the sequence count when placed in low to mid in-degree regions, and also effective in prolonging sequences when mid to high in-degree regions were modulated. Weakening synapses modulated the activity strongest in mid to high in-degree regions. The transmission of STAS through mid in-degree regions was probabilistic (Supplementary Fig. 1), and hence, the most potent modulations were in mid in-degree regions. Major deviation in STAS features (count and duration) points towards a local mechanism that reshapes network dynamics, hence, we further investigated the characteristics of these strong-change locations.

### Motifs underlying the effect of local synaptic changes on STAS dynamics

The observation that a subset of mid in-degree regions were particularly susceptible to perturbations suggested that local synaptic/neural changes might be a mechanism to control STAS dynamics. There are multiple ways in which these local perturbations may arise, for example, from the spatially localized release of neuromodulators (Patriarchi et al. 2018; Glaeser-Khan et al. 2024; Sun et al. 2018), sites of patchy synaptic connections (Lund, Angelucci, and Bressloff 2003), or even volume transmission (Liu, Goel, and Kaeser 2021). In our model, given the geometry of the connectivity, mid in-degree regions could be surrounded by high- and/or low in-degree regions, and that gave rise to specific motifs enabling control over STAS dynamics.

#### Starter

We refer to the mid-degree regions at the start of a pathway as a Start motif (Fig. 3A: left column) because a change in the local connectivity in these regions can affect the activity in two ways: (1) to initiate sequences with a higher probability and (2) to prepend the pathway increasing its overall range.

**Figure 3.**
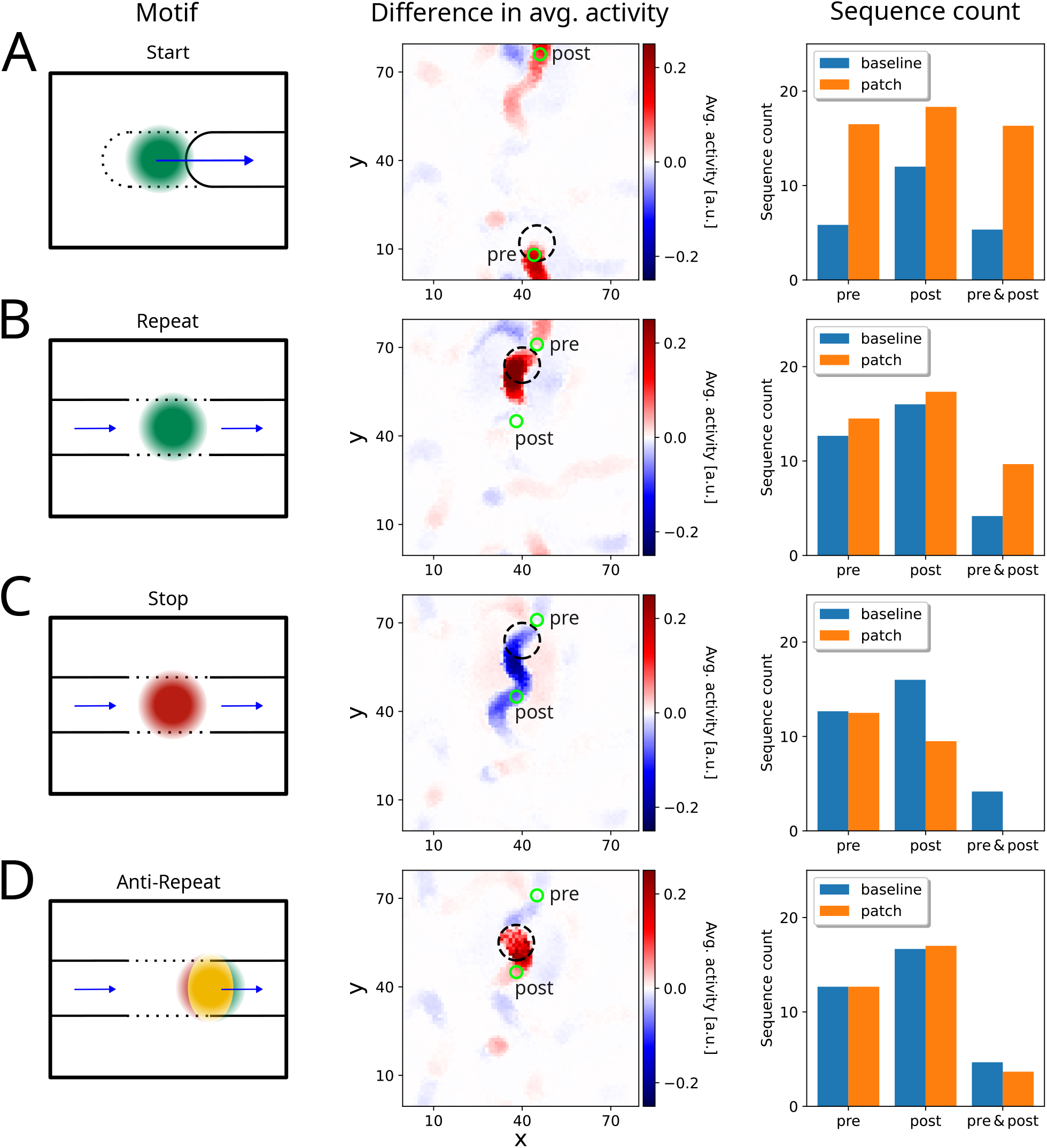
Effect of changes in the local connectivity within a single pathway **(A)** Start motif: (left column): The local patch was at the beginning of a pathway to evoke more sequences and to prepend the pathway. (middle column): Difference between the average activation with a patch compared to baseline activation. (right column): Number of sequences for detection spots *pre, post*, and subsequent activation of *pre & post* (detection spots are indicated by green circles in the middle column). **(B)** Repeat motif: An increase in connectivity of neurons in this location raised the probability of sequence transmission. **(C)** Stop motif: A decrease in connectivity of neurons in this location decreased the probability of sequence transmission. **(D)** Anti-Repeat motif: An increase in the connectivity of neurons in this region triggered the independent emergence of STAS. Such new and spurious STASs competed and could suppress other pre-existing sequences.

To illustrate the effect of Starter locations, we used the same network as shown in Fig. 1D (and its average activity in E). We identified a Starter location (Fig. 3A: middle column, black dashed circle), and studied its effect on the sequence count. For the quantification of local changes in sequence count, we registered all STAS that crossed small spots along the pathway (Fig. 3A: middle column, green circles, for details see Methods).

We found that the Starter patch increased the number of sequences locally, i.e., the spot marked as *pre* (by 184% compared to baseline; Fig. 3A: right column). Given an activation at *pre*, the Starter patch increased the transmission of this activity to the pathway, i.e., the *post* spot (*pre* → *post*: P(*post*|*pre*): 99% (patch) vs 91%(baseline)). Furthermore, in the presence of the Starter patch, if there was an activity at *post*, it was more likely to have originated at *pre* (i.e., P(*pre*|*post*): 89% (patch) vs 44% (baseline)). Thus, the Starter patch increased the transmission along a pathway and also reduced the probability of an independent activation of *post* location by emerging or other (spurious) sequences.

The increased probability of activation in the Starter motif effectively increased the length of the pathways, with more STAS being generated and propagated along the pathway (Fig. 3A: right column). A mid in-degree region could also occur at the end of a pathway, and local changes there also had a similar effect of elongating the pathway. However, such regions did not change the global activity in the network in a measurable manner (not shown).

#### Repeat-Stop

A mid in-degree region may also occur along a pathway, i.e., between two high in-degree regions, featuring unreliable transmission of activity. A decrease in the local connectivity in such regions could essentially ‘stop’ the transmission. Alternatively, an increase in the local connectivity would increase the reliability of the transmission. This is akin to a repeater device in electronic communication systems. Therefore, we refer to these locations as Repeat-Stop motif (Fig. 3B, C: left column).

To illustrate the effect of changing the connectivity strength of the Repeat-Stop motif, we followed the same approach as was used above for the Starter motif (here, the *pre* spot is on the incoming pathway, and the *post* spot is on the outgoing pathway). As expected, increasing the excitability of the Repeat-Stop motif increased the number of sequences that traversed both detection spots *pre & post* indicating a higher transmission probability *pre* → *post* (i.e., P(*post*|*pre*): 76% (patch) vs 37% (baseline)). Note that the overall sequence count at *pre* and *post* did not change much individually, and thus could not account for the increased STAS count of *pre & post* (Fig. 3B: middle and right column). Similarly to the Starter motif, the activation of *post* was shifted towards STAS that already crossed the *pre* location (i.e., P(*pre*|*post*): 59% (patch) vs 28% (baseline)).

By contrast, a decrease in the local connectivity of the Repeat-Stop motif lowered the transmission probability as no sequences traversed both detection spots *pre & post* (i.e., P(*post*|*pre*): 28%; Fig. 3C: right column). In addition, the average activation decreased far downstream along the pathway (Fig. 3C: middle column).

#### Anti-Repeat

It is important to note that the specific location of modulation is crucial for the Repeat-Stop motif. For instance, when we instead increased the local connectivity in a high in-degree region just following a Repeat-Stop motif in a pathway, the previously detected ‘Repeat’ pattern could no longer be observed as the transmission rate *pre* → *post* decreased (i.e., P(*post*|*pre*): 29% (patch) vs 37% (baseline); Fig. 3D). Therefore, we refer to such regions as Anti-Repeat motifs. The increased excitability in the Anti-Repeat motifs evoked sequences independently, and these spurious sequences then suppressed the transmission *pre* → *post* temporarily due to the asymmetric shadowing inhibition (cf. Fig. 1F). Hence, fewer sequences that were detected in *post* originated in *pre* (i.e., P(*pre*|*post*: 22% (patch) vs 28% (baseline); Fig. 3D: right column).

### Mid in-degree regions alter the dynamics of branching and merging of sequences

Thus far, we have discussed motifs that affect a single pathway, affecting their transmission probability and length. Given the geometry of connectivity in our network, pathways merged and split. Here, sequences competed with each other due to the shadowing inhibition. A mid-degree region close to the branching point could bias the activity by modulating the competition or cooperation between pathways. Thus, new motifs such as ‘Gate’ and ‘Select’ arose in the network.

#### Select

Consider a region in the network where one pathway (Main pathway *M*) diverges into two different pathways (Branches *B*_1_, *B*_2_, Fig. 4A-C). Here, it would be advantageous to *select* the branch for further sequence propagation. When the activity from pathway *M* arrives at the branching point, three scenarios could occur: (1) the activity dies as transmission to either branch fails; (2) the activity is transmitted to one of the branches, or (3) both branches become activated. An increased synaptic strength in the Select motif (green patch in Fig. 4A) could bias the activity transmission along one branch (here symbolically to branch *B*_1_), which might also affect activity in the other branch (Fig. 4D-F).

**Figure 4.**
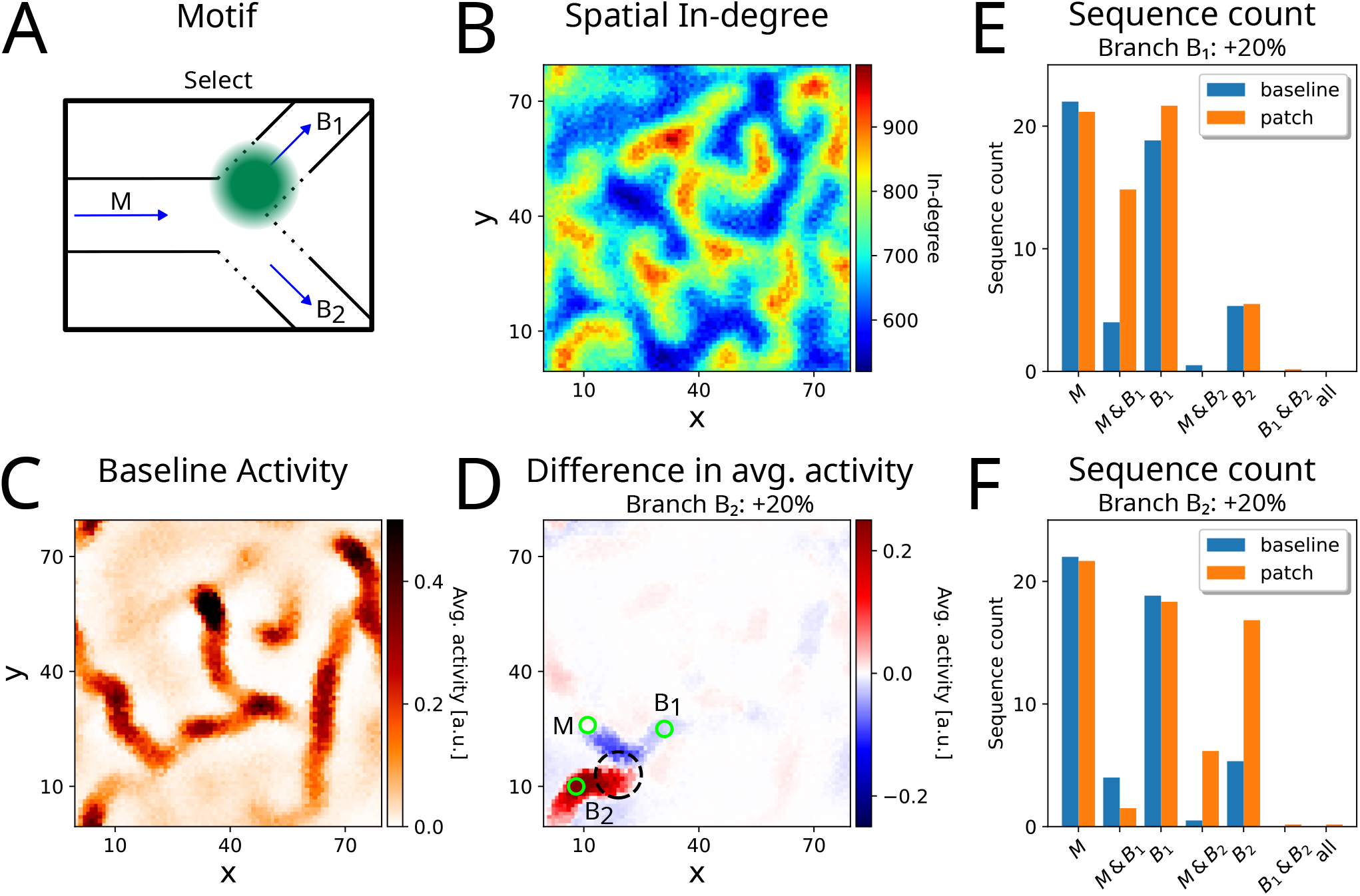
Effect of changes in the local connectivity at a diverging intersection. **(A)** Select motif: A change in the local connectivity of neurons this region could change the probability of selecting one outgoing branch over the other. **(B)** In-degree landscape of the excitatory neurons (see also 1 D). **(C)** Average activity across time. **(D)** Difference in average activity induced by local increased in the connectivity (black dashed circle) and the baseline shown in the panel **C. (E)** Sequence count registered across detection spots on the main pathways *M*, and the branches *B*_1_ and *B*_2_. Strengthening the local connectivity towards branch *B*_1_ increased the transmission rate towards branch *B*_1_. **(F)** Same as in panel **E**, but when local connectivity was increased towards branch *B*_2_.

For instance, modulation of the connectivity towards branch *B*_1_ increased the transmission *M* → *B*_1_ (i.e., P(*B*_1_|*M*): 70% (patch) vs 18% (baseline)) while the *M* → *B*_2_ transmission was slightly reduced (i.e., P(*B*_1_|*M*): 0% (patch) vs 2% (baseline); Fig. 4E). Similarly, we could increase the transmission *M* → *B*_2_ by modulating the connectivity towards branch *B*_2_ (i.e., P(*B*_2_ |*M*): 28% (patch) vs 2% (baseline); P(*B*_1_|*M*): 7% (patch) vs 18% (baseline); Fig. 4F).

As expected, a reduction of local connectivity at the Select motif also changed the dynamics around the intersection. The transmission to the modulated branch *M* → *B*_1_ was essentially blocked (i.e., P(*B*_1_|*M*): 0% (patch) vs 18% (baseline); Supplementary Fig. 2). Interestingly, this kind of perturbation also affected the transmission towards the unmodulated branch. Furthermore, the spontaneous emergence of STAS also changed (sequence count for only *B*_1_ or *B*_2_, respectively). Whether it increased or decreased was dependent on the modulation strength (Supplementary Fig. 2C, D).

#### Gate

In a scenario where two pathways merged, the interaction between two incoming sequences could be modulated by varying the local connectivity in the pathways just before the merging point.

In the simplest case, local weakening of the connectivity of branch *B*_2_ essentially disconnected the *B*_2_ pathway, halting transmission from branch *B*_2_ → *M* (Fig. 5A-C). This effectively *gated* sequence progression, allowing transmission only from the branch *B*_1_ → *M*. Thus, due to the lack of competition of arriving sequences, *B*_1_ → *M* effectively acted as a single pathway (Fig. 5D, E). Consequently, the transmission from *B*_1_ → *M* was increased (i.e., P(*M*|*B*_1_): 71% (patch) vs 31% (baseline)), while transmission probability from *B*_2_ → *M* was abolished (i.e., P(*M*|*B*_2_): 0% (patch) vs 68% (baseline)), hence the name Gate motif.

**Figure 5.**
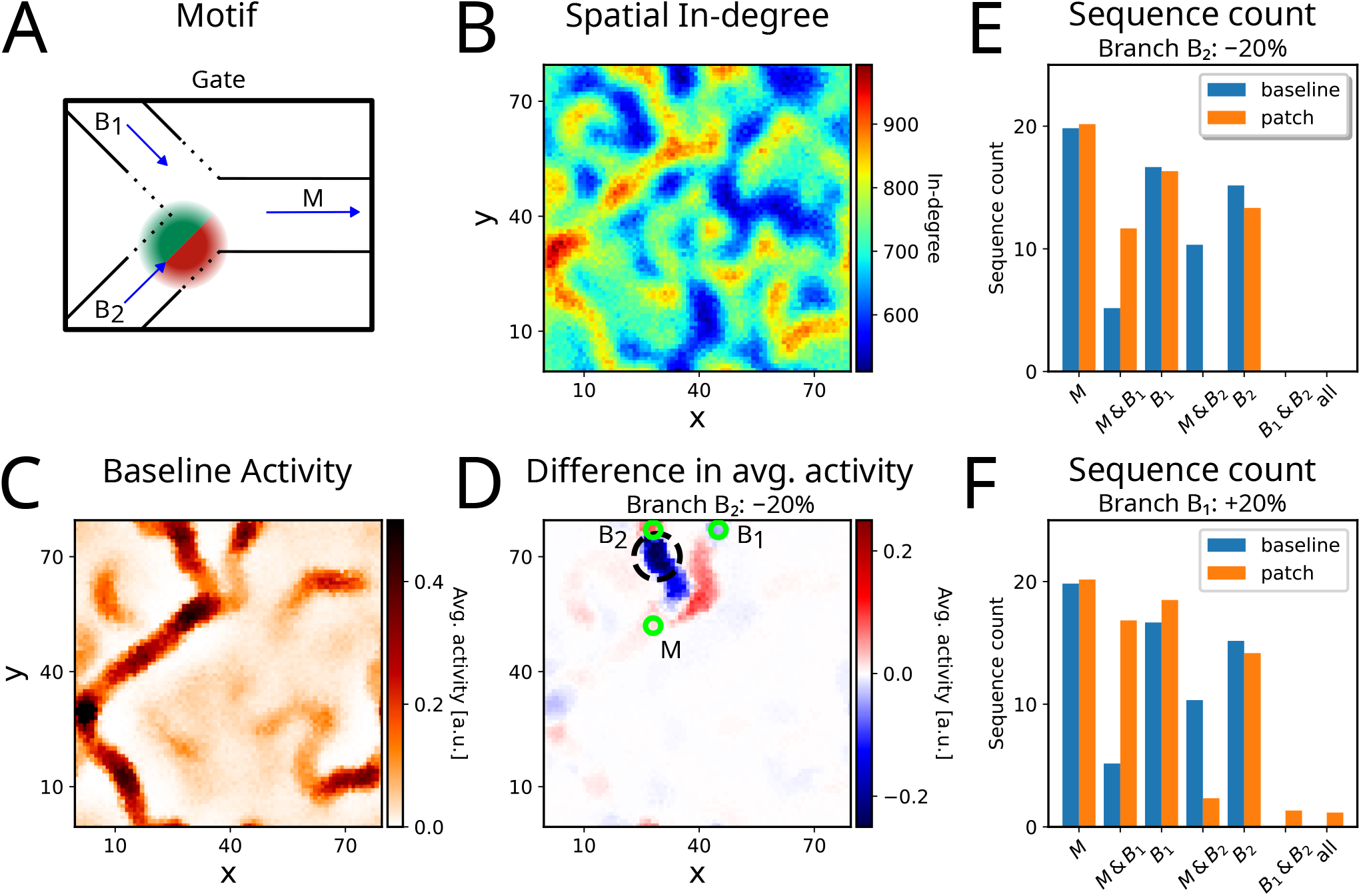
Effect of changes in the local connectivity close to the merging point of two pathways. **(A)**Gate motif: Local modulation on an branch alters the transmission probability towards the main branch. **(B)** In-degree landscape of the excitatory neurons (see also 1 D). **(C)** Average activity across time. **(D)** Difference in average activity when local connectivity in the gate region (black dashed circle) was decreased and baseline shown in the panel **C. (E)** Sequence count registered across detection spots on the branches *B*_1_, and *B*_2_, and the main pathway *M*. A reduction of local connectivity on branch *B*_2_ increases the transmission *B*_1_ → *M*. **(F)** Same as in panel **E** but with an increase of local connectivity on *B*_1_ also resulted in elevated transmission *B*_1_ → *M*.

However, if we strengthened local connectivity in branch *B*_1_, its effect on the transmission unfolded in two ways: In some cases, the elevated activity induced stronger competition between the STAS of *B*_1_ and *B*_2_. As a consequence, the transmission probability from *B*_1_ → *M* was increased (i.e., P(*M*|*B*_1_): 91% (patch) vs 31% (baseline)), while it was reduced from *B*_2_ → *M* (i.e., P(*M*|*B*_2_): 16% (patch) vs 68% (baseline); Fig. 5F). In rare instances, the two incoming STASs excited each other (cooperation) such that both sequences survived and merged despite the recurrent inhibition. Therefore, we also found activation of *all* detection spots: on average, only 2.5 sequences merged per simulation (4 seconds) compared to no observed cooperation in the baseline simulation (Fig. 5F). Interestingly, this effect was more prominent when the modulation was only +10% (Supplementary Fig. 3).

Whether competition or cooperation was emphasized depends on the relative strength and timing of the STASs. Given the connectivity in our network, activity traveling over the excitatory neurons (green region in Fig. 6A, C) created a trailing shadow of inhibition (blue region in Fig. 6A, C) whose strength depended on the size of the excitatory pool recruited by the sequence. Therefore, a strong STAS could destabilize or even suppress a weaker STAS (Fig. 6B). This phenomenon would be especially prominent when one sequence arrives earlier than the other. This asymmetric suppression is also emphasized in the Anti-Repeat motif (see above).

**Figure 6.**
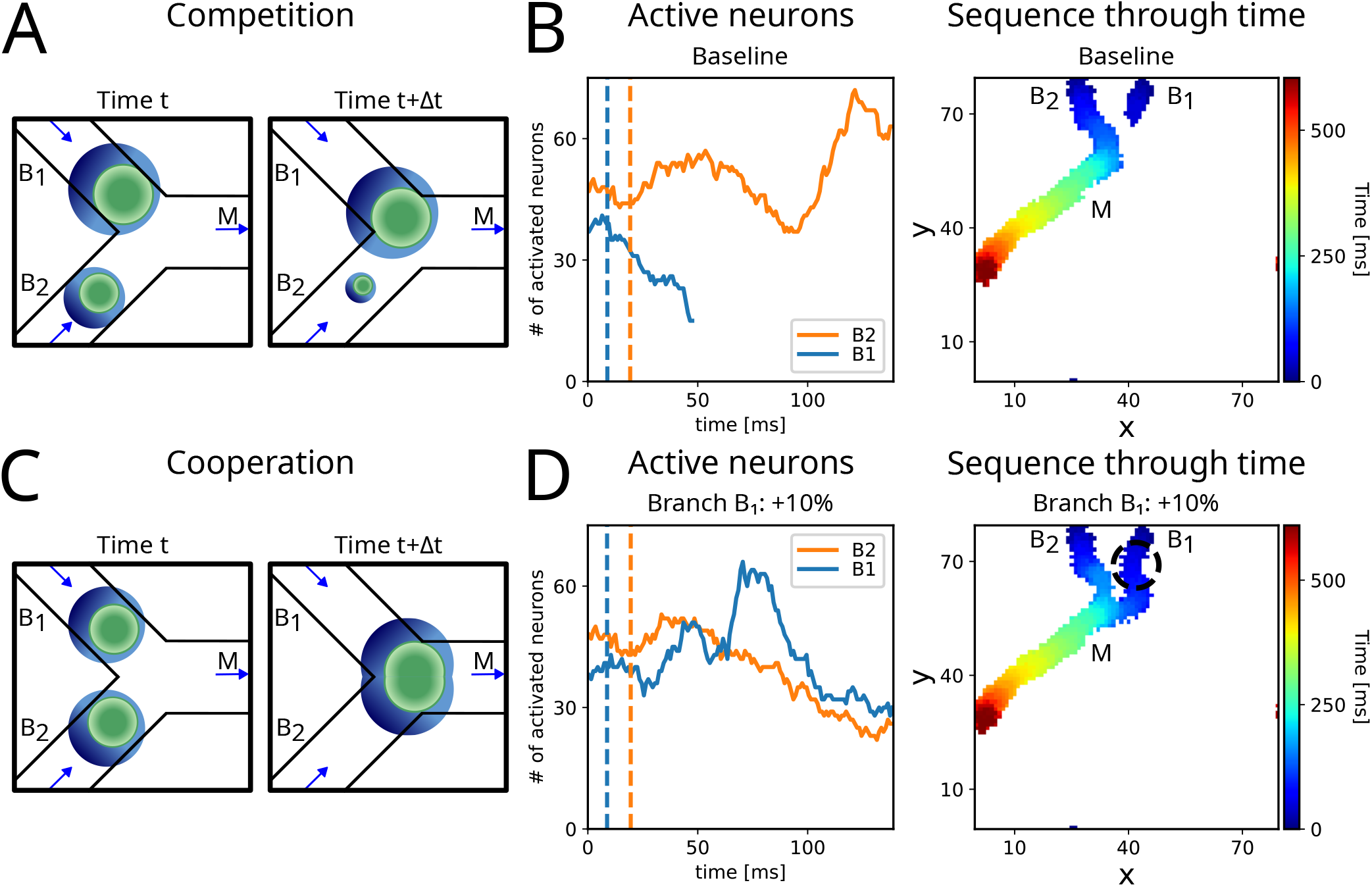
Competition and Cooperation **(A)** Schematic of Competition: When the two incoming sequences are dissimilar in timing or strength, only one lasts, while the second fades. **(B)** (left) Number of active neurons show the balance of the two sequences (on branch *B*_1_ and *B*_2_, respectively). As *B*_2_ was stronger, it suppressed the activity on *B*_1_, and hence, continued to the main branch *M*. Vertical dashed lines indicate the time that the sequences were at the detection spot of the corresponding branch. (right) Activity across space. Color indicate the time point of last activation. **(C)** Schematic of Cooperation: When two balanced sequences arrived simultaneously at the intersection, they cooperated and continued together on the main branch *M*. **(D)** (left) Number of active neurons on each branch (*B*_1_ and *B*_2_) were similar, i.e. balanced before merging. The two sequences merged at *t* = 139ms. (right) Same as in panel B but for Cooperation.

By contrast, if two balanced STASs approached the intersection (almost) simultaneously, the activity could merge, and joint transmission from both branches was possible (Fig. 6D). Merging at an intersection requires recurrent excitation to overcome the surround inhibition, which is weakest at the front of each moving bump of activity. There is thus a window of opportunity for cooperation that depends on the size and relative timing of the two activity sequences. We note that in our networks, cooperation was observed far less frequently than competition.

## Discussion

To explain complex animal behavior, Donald Hebb introduced the idea of ‘Phase Sequences’ (Hebb 1949). Sherrington similarly had envisioned the brain as an ‘enchanted loom’, where “millions of flashing shuttles weave a dissolving pattern, always a meaningful pattern though never an abiding one; a shifting harmony of subpatterns” (Sherrington 1940). These conceptual ideas require that neuronal networks in the brain can not only exhibit sequential activity but also rapidly switch and reconfigure the sequences. A number of mechanisms exist to explain the emergence sequences (e.g., Hebbian plasticity, symmetry breaking due to synaptic depression or neuron adaptation, and asymmetric connectivity due to neuron morphologies). However, the problem of rapid switching among different sequences has received little attention. In some sense, this question of rapid switching relates to a question Larry Abbott posed ‘Where are the switches on this thing?’ (L. Abbott 2006).

### Small switches with big leverage

In a network that is intrinsically poised to generate spatio-temporal sequences, ‘switches’ are needed to control the magnitude (on/off/amplify) and direction of the activity. A solution to this problem is to control the magnitude (Vogels and L. Abbott 2009) or timing of excitation and inhibition (Kremkow, Aertsen, and Kumar 2010). Here we show that in networks with spatially correlated anisotropic connectivity of individual neurons (Spreizer, Aertsen, and Kumar 2019) several locations naturally emerge that can guide sequential activity. A small (±20%) modulation of connectivity of as few as 50 neurons in these regions is sufficient to flexibly route sequential activity.

These observations suggest that stimulation of heterogeneous networks in the brain should be location-specific. In fact, there should be regions where activation/inactivation of neurons will elicit a rich network response, while other locations will display no visible impact on the network activity. The magnitude of the network response will, however, also depends on the strength of the modulation (cf. Supplementary Fig. 2, 3)

Moreover, spatially localized stimulation is more likely to generate a larger response than stimulation of neurons randomly distributed in the network. Note that here a large response implies that we should observe large reshuffling of activity at the population level as sequential activity would be rerouted – recruiting new neurons and silencing others (see Fig. 2). Thus a local modulation of such ‘switches’ could effectively alter the dimensionality and geometry of the manifold of neural activity.

### Where are the locations that have a big impact on network activity?

Neuron degree (in- or out-degree) in the brain is likely to be heterogeneous, given the neuronal chemical specificity and morphology (i.e., axonal and dendritic arbors are not isotropic in space). Furthermore, synaptic plasticity may also introduce additional heterogeneity in network connectivity. Thus, some neurons will have more connections than others. When the heterogeneity of neuron degree is uniformly distributed in the space, it may not have much impact (Spreizer, Aertsen, and Kumar 2019). However, when there are regions of high, medium, or low degree, non-trivial activity dynamics emerge (Spreizer, Aertsen, and Kumar 2019). From the perspective of controlling the activity dynamics, interactions among different degree regions play a crucial role.

Low in-degree neurons do not see much of the network activity, and therefore changing their connectivity strength by a small amount is not very effective in influencing the network dynamics. On the other hand, high in-degree regions already receive strong input from the network, and therefore small change in their connectivity also does not have much influence on the network activity. By contrast, medium in-degree regions are well poised to take advantage of the small modifications in connectivity/excitability and switch the network activity in a big way. The impact of a change in the connectivity/excitability of such neurons depends on the local neighborhood. Various motifs arise that can supervise sequence transmission and steer sequential activity at pathway intersections (Fig. 3, 4, 5).

In our model, the spatial distribution of neuron in-degree determines their impact on the network dynamics. To some extent, a distribution of synaptic strengths can also do the same. Recently, Riquelme et al. (2023) showed that in a network, when synaptic weights are drawn from a log-normal distribution, feedforward pathways emerge where neurons are connected by strong synapses. In that model, feedforward pathways are linked by weaker connections, and stimulation of these links grants effective control of the sequential activity. Since connections in the mammalian cortex are relatively weaker, our mechanism may be more suited for the control of activity in the mammalian brain. Here, we have argued that certain medium in-degree neurons form the basis of high-impact locations. This is simply because in our model in-degree is variable. The argument should also hold when either out-degree or both in- and out-degree are heterogeneous and have a spatial distribution, but this remains to be shown.

### Targeting the high-impact locations

How could upstream networks target these neurons (spatial locations in the network) that have a high impact on network activity? One possibility is that projections from an upstream network impinge in a spatially compact manner. This external input can directly drive the neurons closer to the threshold. This input, of course, has to be precisely timed with the arrival of a sequence to enable (or disable) further propagation. Moreover, such a stimulus could be strong enough to inject an additional sequence into the network.

Alternatively, neuron excitability or connectivity can be transiently modulated by neuromodulators which are released in a spatially patchy manner (Patriarchi et al. 2018; Hamid, Frank, and Moore 2021; Sun et al. 2018; Glaeser-Khan et al. 2024). Neuromodulation-induced changes in the excitability or synaptic strength can occur on behaviorally relevant time scales (hundreds of ms). Neuromodulatory neurons typically have a rather large axonal arbor, and they can simultaneously target neurons with different in-degrees in a non-specific manner. However, the effect of neuromodulatory action may be manifested via the neurons with medium in-degree.

Finally, synaptic plasticity and learning could also modify the connectivity of incoming projections over longer time scales and bias it towards medium in-degree neurons. A correlation-based learning rule could unfold the synaptic weights in the following way: due to the high intrinsic activity of pathways, the correlation of input and activity would be low. On the other extreme, low in-degree regions are mostly silent and hence would not follow the input drive either. However, a medium in-degree region would be highly responsive to the input and thus form the strongest synapses. Consequently, inputs from an upstream network may be systematically potentiated if they impinge on medium in-degree neurons.

### What does it mean for computations in the brain?

Performing any task in a natural environment involves multiple steps. If each component of a task (e.g., extending the arm, opening the finger, etc.) is represented by sequential activity of neurons, then behavior should involve dynamic interactions among sequences. Cooperation and competition naturally arise, but strengthening, stopping, gating, and routing should be possible. The ‘switches’ we have described can form an important component of neural circuitry that allows for dynamic control of sequences.

In a cognitive or sensorimotor task, input may initiate a sequence (for instance, seq#1 in Fig. 7). Depending on how the ‘switches’ (blue start/repeat/select/gate) are activated by modulatory or contextual input, the sequence may traverse the network and reach seq#A or seq#B (representing a conceptual read-out). In a large network with many pathways and multiple motifs simultaneously, sequences may start in one pathway and dynamically choose their trajectory in the network (Fig. 7). That is, a sequence originating from one pathway can arrive at various endpoints, but also one endpoint can be reached by many initial pathways.

**Figure 7.**
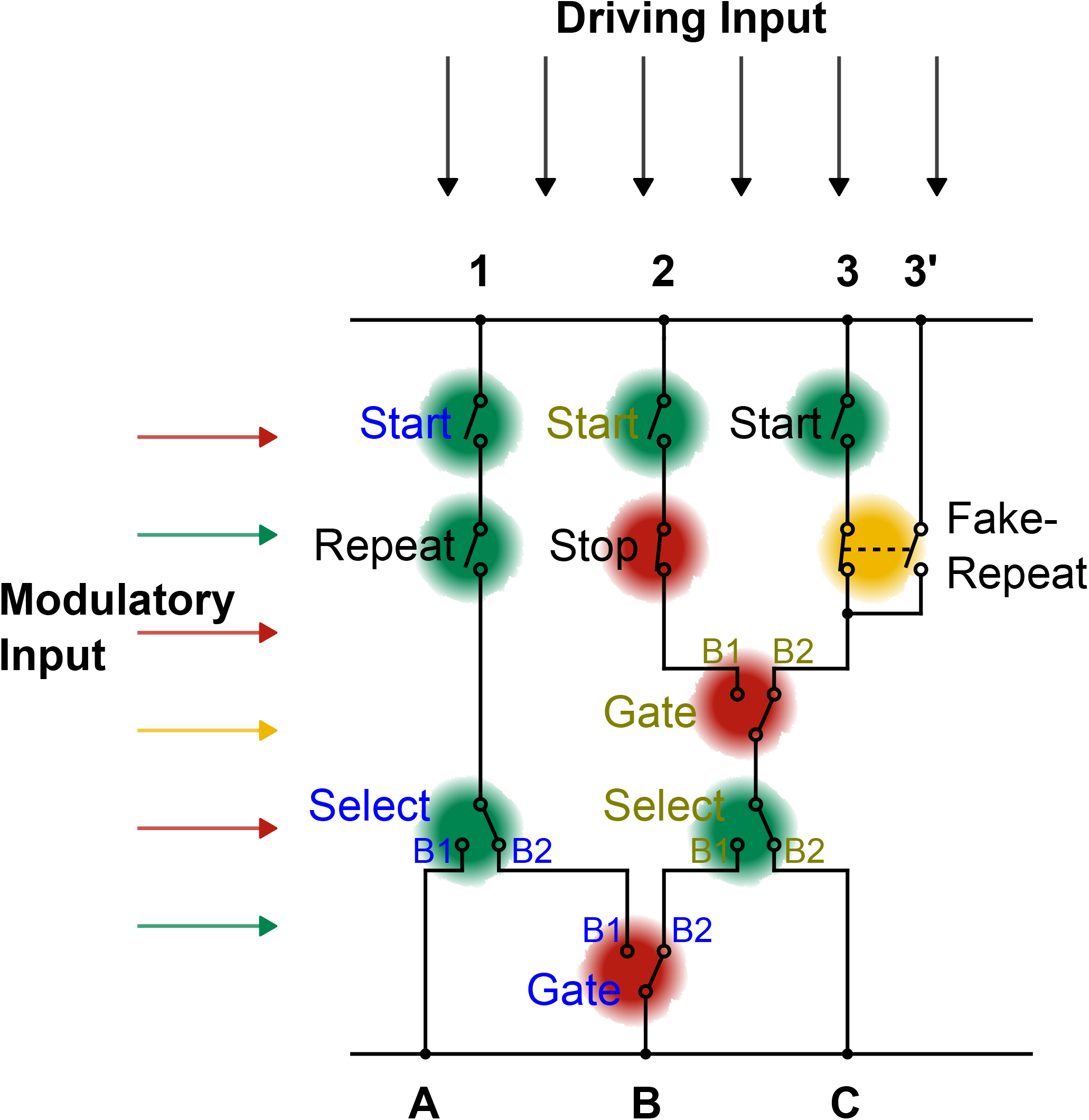
Schematic of the neural circuit. The driving input ignites sequences at the entry points 1, 2, 3, which can be modulated by the corresponding Start-motif. The modulatory input sets the switches, which results in various possible trajectories the sequence traverses. The read-out A, B, C can then be reached by sequences originated from multiple entry points.

This multiplexing creates a framework for fast and flexible computations, while the individual parts of the pathways remain rigid and reliable. Other properties of sequences, namely the number of neurons recruited at any given point in time and the propagation velocity, may also play a role and contribute to the computational power (Lehr, Kumar, and Tetzlaff 2023; Lehr, Kumar, and Tetzlaff 2024). Although not investigated in this study, background activity or forms of short-term synaptic plasticity may regulate sequence dynamics and propagation velocity (Murray and Escola 2017).

### Model Predictions

Despite simplifications and assumptions (as needed by models), our model makes several testable predictions that could provide data to both support the model and to extend it. First and foremost, stimulus response should depend on spatial locations. That is, local stimulation will evoke richer network responses compared to spatially distributed input, as shown in Fig. 2. In terms of network connectivity, projections from an upstream network should be spatially compact and should target neurons that have mid-range in-degree in the local network.

Given the presence of ‘switches’, sequences traversing only parts of a pathway would be highly reliable, and the full sequence is finally composed of a few (or many) shorter elemental sequences. This perspective interprets trial-by-trial variability as a flexible web of computations.

In most brain regions, multiple neuromodulators converge simultaneously in space and/or time. In our study, such convergence of antagonistic (i.e. one NM excites and the other inhibits) neuromodulators on the same neurons can provide bidirectional control of the ‘switches’. In fact, regions where multiple neuromodulators converge could be the ‘switches’ of the network.

## Material & Methods

### Neuron model

Here each neuron was modeled a firing -rate based unit. The firing rate dynamics were modeled as:

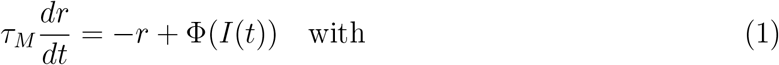

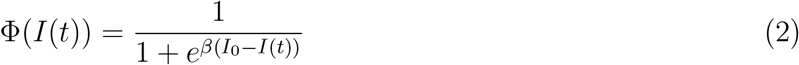

where *τ*_*M*_ is the membrane time constant. *β* determines the steepness of the transfer function, and *I*_0_ defines the external input that results in an output of *r*_0_ = 0.5 in the absence of any network connectivity.

The total input to a postsynaptic neuron is defined as

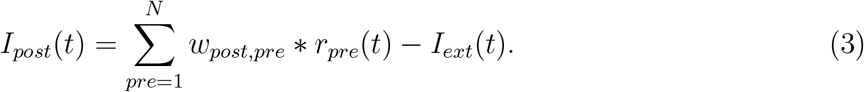

The weights *w* and the external input *I*_*ext*_ will be discussed in detail later. All parameters of the network and neurons are given in 1.

### Network model

The network consisted of 6400 excitatory neurons and 1600 inhibitory neurons, called excitatory and inhibitory population, respectively. This resembles a ratio of 4:1 as found in the mouse neocortex (Braitenberg and Schüz 1998). The neurons were placed on an 80-by-80 grid with equal spacing. The boundaries of the grid were forded to form a torus. This was done to avoid boundary effects. The inhibitory neurons were places at the center of four excitatory neurons, ensuring equidistance among them (Fig. 1A).

Each neuron formed 2400 (600) connections to the excitatory (inhibitory) population. Multiple connections were allowed, while self-connections were prohibited. The connection probability followed a Gaussian distribution, with different spread *σ*_*p*_ for the excitatory and the inhibitory population,

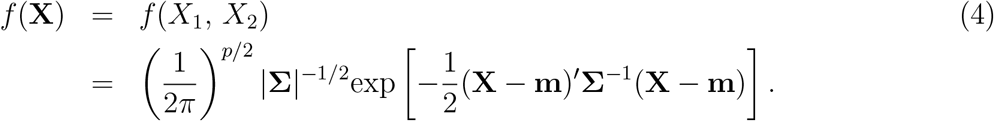

**Table 1.**
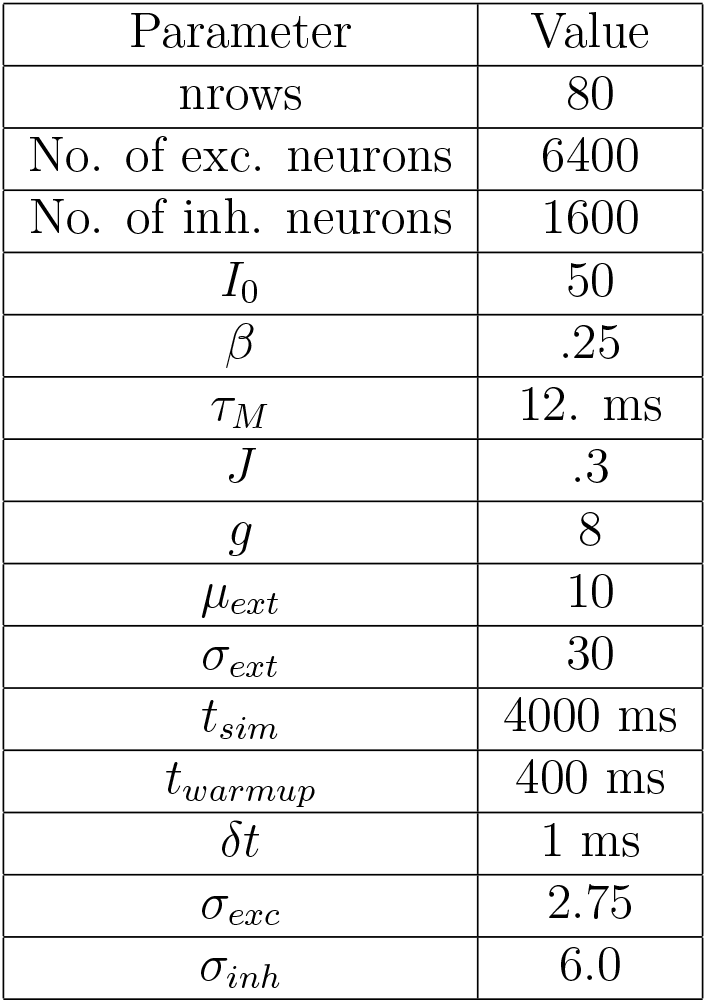
Parameter of the network and its neurons.

| Parameter | Value |
| --- | --- |
| nrows | 80 |
| No. of exc. neurons | 6400 |
| No. of inh. neurons | 1600 |
| $I_0$ | 50 |
| $\beta$ | .25 |
| $\tau_M$ | 12. ms |
| $J$ | .3 |
| $g$ | 8 |
| $\mu_{ext}$ | 10 |
| $\sigma_{ext}$ | 30 |
| $t_{sim}$ | 4000 ms |
| $t_{warmup}$ | 400 ms |
| $\delta t$ | 1 ms |
| $\sigma_{exc}$ | 2.75 |
| $\sigma_{inh}$ | 6.0 |

In this 2D case with no correlation among the directions, the width of the distributions was

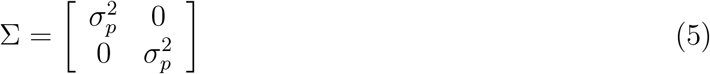

with *pϵ*{exc, inh} for the excitatory and inhibitory population, respectively. The higher spatial spread of the inhibitory population resulted in an effectively Mexican-hat connectivity kernel with local excitation, but long-range inhibition (Fig. 1A).

The overall connection strength *w*_*post,pre*_ between the *pre*synaptic and the *post* synaptic was calculates as

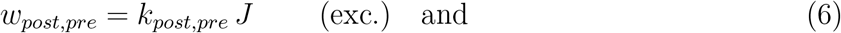

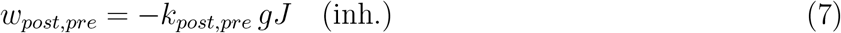

with *k*_*post,pre*_ being the number of formed connections, *J* was taken as the synaptic weight, and *g* the ratio of inhibitory and excitatory synapses.

### Shifts/Perlin landscape

As Spreizer, Aertsen, and Kumar (2019) showed, spatio-temporal activity sequences arise when (1) individual neurons preferentially make some their connections in a specific direction *ϕ*, and(2) the *ϕ*’s of neighboring neurons are similar. The spatial correlations in the network are based on a 2-D Simplex noise - a class of gradient noise (Perlin 2001). The spatial extent of the correlation is controlled by the *size* parameter and relates loosely to the width of a pathway. Each 2-D sample results in a different landscape, thus leading to various spatial motifs, which are discussed later.

### External input

Additional external current was modeled as an approximation of Gaussian white noise (GWN), with mean *µ*_*ext*_ and variance *σ*_*ext*_. The external input was sampled independently for each neuron.

### Network simulation

The network activity was simulated for *t*_*sim*_ = 4000ms, with a preceding warm-up phase (*t*_*warmup*_ = 400ms). As different sources of stochasticity exist in the model, we ran the simulations re-sampling the external noise input each time. For simplicity, the connectivity was fixed across the implementations of the GWN. We tested that the dynamics were robust against randomness in the connectivity. Thus, we defined the baseline activity across a connectivity landscape for the simulation time *t*_*sim*_, and with an implementation/seed of the external noise leading to *S*_*seeds*_ baseline simulation.

The network was simulated using the brian2 framework (Stimberg, Brette, and Goodman 2019).

## Neuromodulation

To modulate the sequence dynamics, we defined a small local neighborhood called a patch, with a center *c*, and radius *r* (*r* = 6 grid points). From each patch, we randomly chose *N*_*rec*_ = 50 neurons. For these neurons, all incoming synapses were modulated - i.e., the weights were either increased or decreased by *p* = ±20% (or *p* = +10% for one motif). This kind of modulation of local connectivity is inspired by the way neuromodulators act in the brainNadim and Bucher 2014; Patriarchi et al. 2018. Therefore, with the assumption that neuromodulators often act on a longer timescale than neural activity, the patch was active throughout a simulation i.e., the connection strength remained higher or lower through a specific run of the network.

## Detection of sequences

A sequential activation of neurons across space and time was detected as sequences following a multistep algorithm. First, the rate of each neuron was translated into pseudo-spikes by thresholding (Θ = 0.3). These spikes were then clustered using the Density-Based Spatial Clustering of Applications with Noise algorithm (DBSCAN)(Ester et al. 1996) to remove spikes that were not participating in a sequence (*ϵ* = 5, *min samples* = 30, (Pedregosa et al. 2011)). Note that the dimensions of the data were in x-, y-direction, and across time. Although rescaling of the time axis could change the relation of space in time, it appeared not to have a strong effect.

As the network space was a torus, the standard DBSCAN cannot account for activity that convolves around the edges. To account for this, the activity was shifted by *nrows/*2 in x- and y-directions. Then, the DBSCAN clustered the spikes again. Afterward, merging the clusters of both runs, traversing sequences were identified.

Finally, the number of sequences was determined in a general or a location-specific way. Generally, the number of detected clusters also led to the number of sequences in the network. Moreover, every active neuron in any sequence incremented the sequence count for the corresponding coordinate (see Supplementary Fig. 1). When investigating sequences in specific locations, the sequence count was incremented whenever any neuron at a detection spot (circle with radius *r* = 2) participated in a cluster. That way, variability across activated neurons was removed. Hence, it was the sequence count on a pathway instead of the activations of individual neurons in a sequence.

The duration of a sequence was derived from the first and last time points at which the sequence was registered.

## Code availability

The code for reproducing the simulations is made available upon request.

## Author contributions

Hauke O. Wernecke, Andrew B. Lehr, and Arvind Kumar designed the study and interpreted the data; H.W. analyzed the data, developed code, and performed simulations; H.W. wrote the manuscript with contributions from all co-authors.

## 1 Supplementary Figures

**Supplementary Figure 1.**
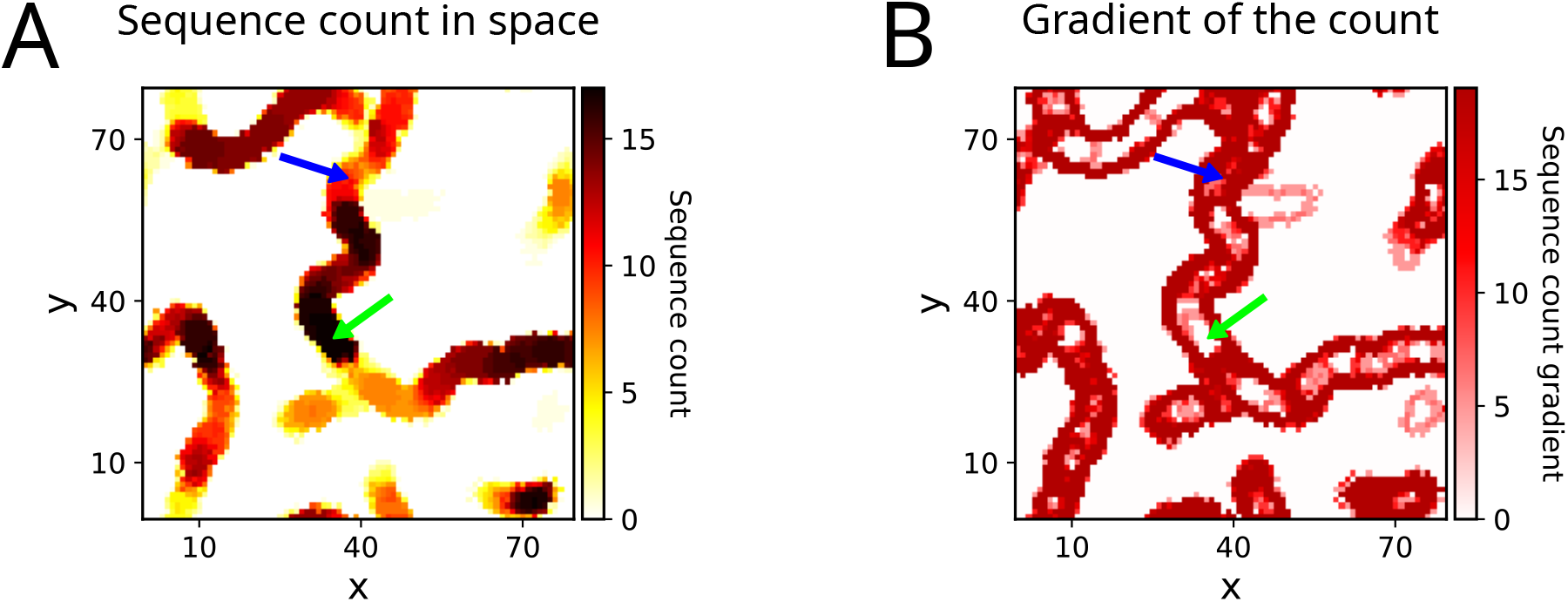
Sequence count and its gradient with respect to space in a across baseline simulations. **(A)** Sequence count is indicated by darker colors. **(B)** Reliability is high if the sequence count was high and the gradient was low, e.g., green arrow. By contrast, unreliable transmission was indicated by a higher gradient within pathways (i.e., not the edges to regions of zero sequence count), e.g., blue arrow.

**Supplementary Figure 2.**
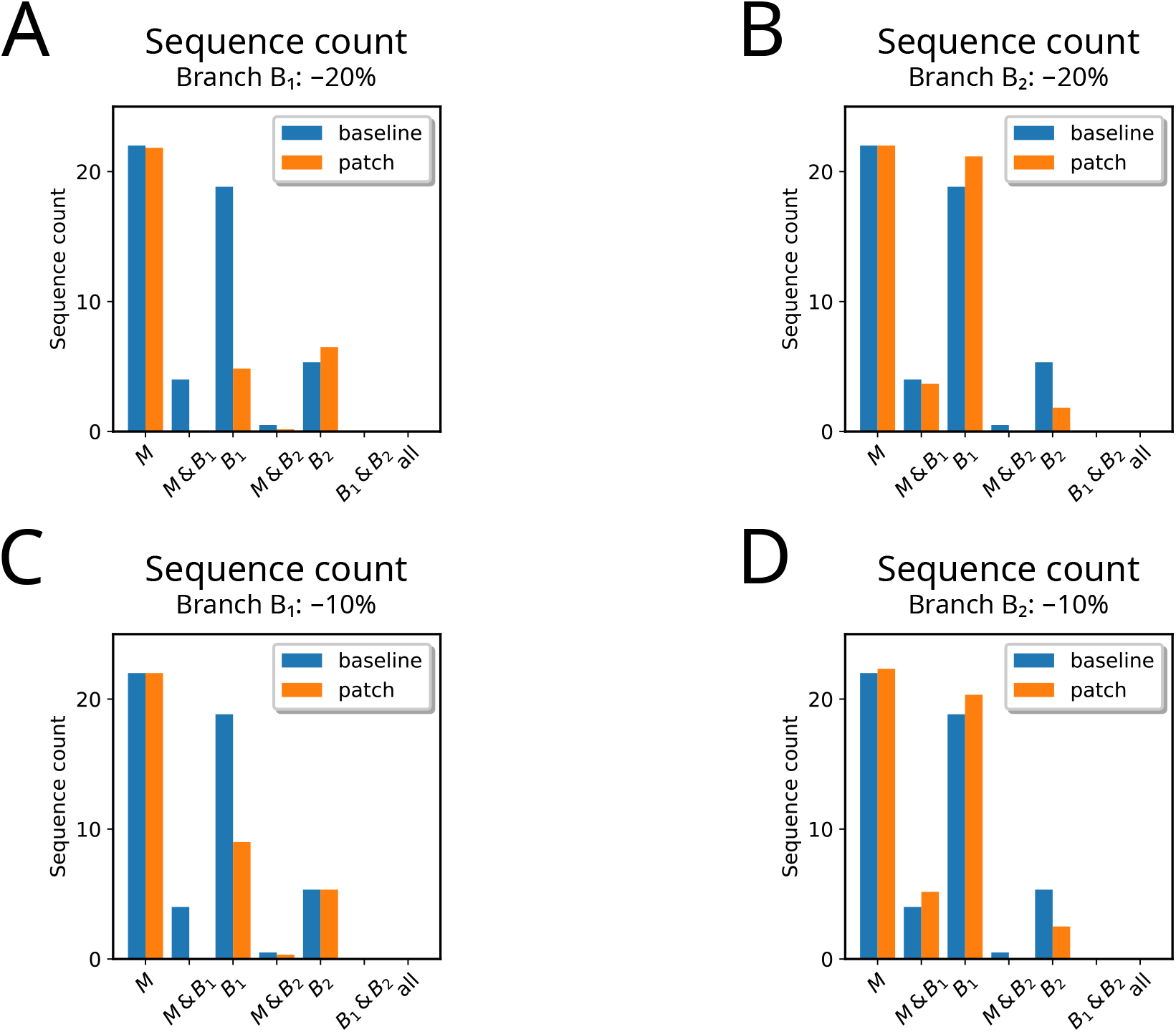
A decrease in the local connectivity effectively acted as a Stop-motif. The transmission to the unmodulated branch might be unaffected (A), or even leveraged due to reduced competition (B).

**Supplementary Figure 3.**
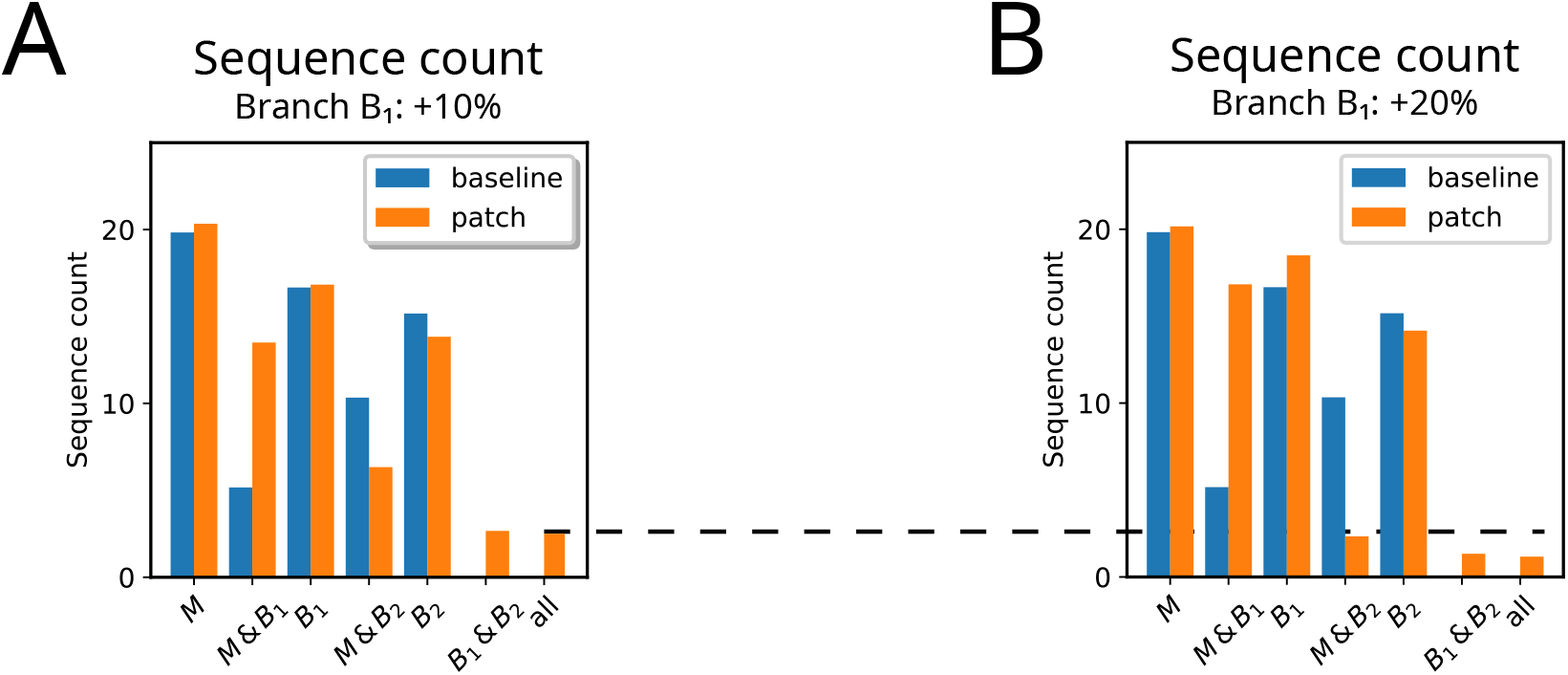
Cooperation was observed more frequently when the modulation strength was only +10% (A), opposed to the standard +20% (B) (cf. dashed line). Sequences detected across *all* detection spots: 2.5 (A) vs 1.2 (B).

